# Genomic signatures of selection in the mound-building mouse *Mus spicilegus*

**DOI:** 10.64898/2026.08.21.746306

**Authors:** Mara C. Baylis, Kayleigh Fort, Samantha Jackson, Linda Wilbrecht, Rachel B. Brem

## Abstract

The steppe mouse *Mus spicilegus* is distinguished by several unusual phenotypes: animals of this species build complex mounds from brush and dirt, and this behavior as well as the pace of development is regulated by season of birth. The genetic and evolutionary basis of species-defining traits in *Mus spicilegus* is unknown. To discover genomic clues to the mechanisms at play, we took a molecular-evolution approach. Sequence-based tests of protein-coding exons, and of brain-active *cis*-regulatory regions, revealed a striking excess of evidence for evolutionary acceleration in the *M. spicilegus* genome relative to those of closely related species. Transcriptional analyses also revealed unique expression programs in the *M. spicilegus* brain, which could be traced in part to *cis*-regulatory sequence variation. Hit loci from our molecular-evolution tests tended to be active in the cerebral cortices and hypothalamus and have annotated functions in neuronal development, neural plasticity, or endocrine signaling. In many cases, their orthologs in other animal systems had been implicated in neoteny/the pace of maturation, unusual building behaviors, and autism. Together, our data indicate that *M. spicilegus* has undergone divergence in brain-active genes to an extent far more than its relatives. We interpret the affected genes under a model in which they mediate the seasonal capacity by *M. spicilegus* for protracted maturation and mound-building. Thus our work provides an evolutionary interpretation of, and catalog of candidate determinants for, shifts in motivated behavior and development in this intriguing mouse species.

## Introduction

Understanding how genomes evolve to establish novel behavioral innovations is a central challenge in comparative neuroscience. As a field, we can identify genes essential for neuronal development or hormonal signaling, but we cannot yet look at a genome and predict how evolutionary change in its sequence or regulation will modulate an organism’s behavior. The field has advanced most in fruit flies, nematodes, and fish, where decades of work tie specific neurons and genes to naturally varying behaviors such as aggression, foraging, olfaction, and circadian rhythms (Fages et al. 2026; Ding et al. 2016; Bertolini et al. 2026; Jernigan et al. 2023; Kent et al. 2009; Sawyer et al. 1997; Shorter et al. 2015; de Bono and Bargmann 1998; Kowalko et al. 2013; Peichel et al. 2001; Palavicino-Maggio and Sengupta 2022). In mammals, in landmark cases it has been possible to establish causal links from sequence to circuit to behavior (Christmas et al. 2023; Lim et al. 2004; Insel and Shapiro 1992; Winslow et al. 1993; Bendesky et al. 2017; Kingsley et al. 2024; Baier et al. 2025; Hu and Hoekstra 2017), but behavioral changes in domesticated laboratory strains, and the paucity of tractable wild models, have limited progress in the field.

The steppe mouse or mound-building mouse, *Mus spicilegus*, is well-suited for the study of novel complex behaviors as well as the regulation of the pace of developmental milestones. Developmental life history in *M. spicilegus* is seasonal: juveniles born in late summer and autumn build vegetation-filled mounds over burrows which they inhabit in groups for the winter; these animals delay dispersal and reproduction until the following spring (Gouat et al. 2003a; Tong and Hoekstra 2012). By contrast, juveniles born in spring do not build mounds and disperse and reproduce within the summer months (Poteaux et al. 2008; Gouat et al. 2003a). Laboratory experiments indicate that social cues and photoperiod mediate seasonal gating of *M. spicilegus* development and reproduction (Simeonovska-Nikolova and Mehmed 2009; Lafaille and Féron 2014; Bárdos et al. 2024; Cryns et al. 2022a; Groó et al. 2013; Bardet et al. 2007).

The evolutionary and genetic basis of *M. spicilegus*’s unusual traits is unknown. Genomic methods have great potential to fill the knowledge gap, since *M. spicilegus* diverged from its close relatives relatively recently (on the order of ∼1 MYA), providing a window in which lineage-specific changes are likely to be interpretable and not overwhelmed by deep divergence (Suzuki et al. 2004). We hypothesized that the unusual phenotypes of interest in *M. spicilegus*, including behavior, were an advantage for the organism in its niche and had evolved under positive selection during its divergence from the rest of *Mus*. We set out to test this notion with a molecular-evolution approach applied to observations of sequence and expression across the genus. Our goal was to trace patterns of non-neutral molecular variation to provide broad insights into the evolutionary history of *M. spicilegus*, and to interpret loci with signatures of selection as candidate determinants of adaptive traits in this system.

## Results

### Amino-acid variation driven by positive selection in *M. spicilegus*

To identify derived alleles in *M. spicilegus* following patterns consistent with a history of positive selection, we first focused on changes in amino-acid sequence, using codon branch models implemented in PAML (Z. Yang 1997; 2007). For each gene, we assessed the evolutionary forces that had driven protein evolution in the type strain of each *Mus* species in turn, and we tabulated the genes emerging as hits in each case. In the results, the detection rate for evidence of adaptive protein evolution was ∼4-fold higher in *M. spicilegus* than in any other species, with 44 loci called as hits (Figure 1a and Supplementary Table 1). These inferences of positive selection in *M. spicilegus* included many implicated in hypothalamic functions, including the regulation of pubertal maturation (*Tmem108)* (Rezende et al. 2023; Akram et al. 2023), body temperature (*Gabre,* the epsilon subunit of the GABA receptor) (Wang et al. 2024), water consumption (*Pcsk1n*) (Aryal et al. 2022), feeding and body weight (*Sdc3*) (Reizes et al. 2001), and the circadian clock (*Per3*) (Matsumura and Akashi 2017). We also noted, among loci with adaptive protein evolution in *M. spicilegus,* genes functioning elsewhere in the brain*, e.g. Vglut2,* a subcortical glutamate transporter (Kaneko and Fujiyama 2002), and *Sytl4*, which is expressed in the bed nucleus of the stria terminalis and implicated in male mating (X. Xu et al. 2012). Another hit locus, *Zmpste24,* is implicated in nuclear envelope function and mediates accelerated aging when knocked out (Varela et al. 2005).

**Figure 1.**
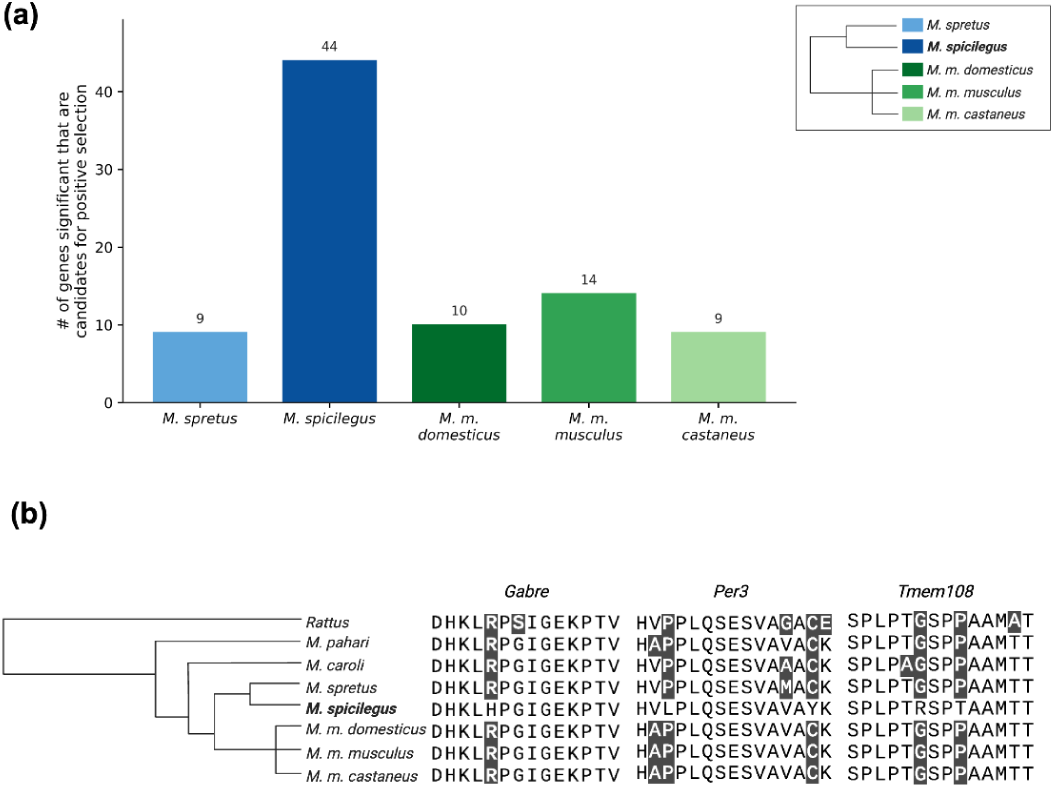
Elevated protein evolution in *M. spicilegus.* Shown are results from tests for adaptive protein evolution across *Mus* via the branch model in the PAML package. (a) Each color reports results of tests for selection on the indicated *Mus* branch. The *y*-axis reports the number of genes with significant evidence for positive selection after multiple-testing correction; gene hit counts are shown above each bar. (b) Examples of loci with inference of adaptive protein evolution in *M. spicilegus*. Shading reports lineage-specific amino-acid substitutions. *Gabre* activity modulates GnRH neuron responsiveness, influencing the timing of puberty and response to sex steroids. *Per3* is a core component of the circadian clock system, primarily regulating sleep-wake cycles, sleep homeostasis, and behavioral responses to light. *Tmem108* acts in neuronal development and synaptic function, promotes dendrite outgrowth and excitatory synaptic activity, and is implicated in pubertal delay (Z. Wang et al. 2024; Matsumura and Akashi 2017; Jiao et al. 2017). Alt text: Two-panel figure showing lineage specific protein coding evolution in *Mus*. A bar chart shows substantially more genes with significant lineage-specific evolutionary rate shifts in *M. spicilegus* than in the other *Mus* lineages tested. A phylogeny and representative protein sequence alignments highlight *M. spicilegus* specific amino acid differences in *Gabre, Per3,* and *Tmem108*.

To explore more systematically the potential function in the brain of loci with accelerated amino acid variation in *M. spicilegus*, we turned to an unbiased analysis of their cell-type-specific expression. We first mapped each gene in the genome to the cell type in which it was expressed highest in brains of two-month-old lab mice, from the Allen Mouse Brain Atlas (Allen Brain Atlas 2026). We then tested whether a given cell type had an elevated count of such peaks among the hits from our PAML screen of the *M. spicilegus* proteome, relative to a random expectation. The results revealed enrichment for expression of our focal genes in the median eminence, arcuate hypothalamic nucleus, and other regions of the hypothalamus, driven by *Gabre, Vglut2*, *Sytl4, Per3, Sdc3* and *Zmpste24* (Supplementary Figure 1 and Supplementary Table 2). We conclude that the *M. spicilegus* proteome has undergone extensive change as the species diverged from its relatives across *Mus*, including at hypothalamic genes with potential relevance to the capacity for delayed maturation and reproduction that are expressed when this species is born in autumn.

### Accelerated sequence evolution in brain-active *cis*-regulatory regions in *M. spicilegus*

For a complementary study of evidence for natural selection in the *M. spicilegus* genome that could shed light on drivers of trait innovations in this species, we turned next to analyses of *cis-*regulatory sequence. Our approach used the phyloP tool, which evaluates sequence variation on a phylogeny for evidence of acceleration or conservation relative to a neutral model (Pollard et al. 2010). We hypothesized that regulation in the *M. spicilegus* nervous system could have been a primary target of evolutionary remodeling. To explore this, we first used as input for our phylogenetic analyses the *Mus* type strain sequences at a set of 246,543 candidate *cis*-regulatory elements (cCREs) identified as DNase accessible sites in assays of whole brain from laboratory mice at two months of age (Pollard et al. 2010; Moore et al. 2026). Across which, we detected 273 loci with evidence for accelerated sequence variation in *M. spicilegus,* ∼10-fold more than in any other *Mus* (Figure 2a and Supplementary Table 3a), dovetailing with the detection rates we had detected in our analyses of protein variation (Figure 1). Controls in which we eliminated *M. spicilegus* sequence from the input data yielded similar hit rates for other species in the set at whole brain-active cCREs, ruling out artifacts from *M. spicilegus* on detection of evolutionary signal in the rest of the tree (Supplementary Figure 2 and Supplementary Table 3b). To assess the extent of accelerated sequence evolution genome-wide, we repeated our analyses of the full phylogeny on cCREs active in heart, liver, and kidney (Sanchez-Contreras et al. 2023; Zhang et al. 2024), finding no marked difference in phyloP detection rate between *M. spicilegus* and other species (Figure 2a). These data make clear that non-neutral sequence divergence is particularly prevalent in the *M. spicilegus* genome at regulatory loci active in the brain, but not in other tissues, as expected if the resulting variants contributed to this species’ motivated behavior and/or development traits.

**Figure 2.**
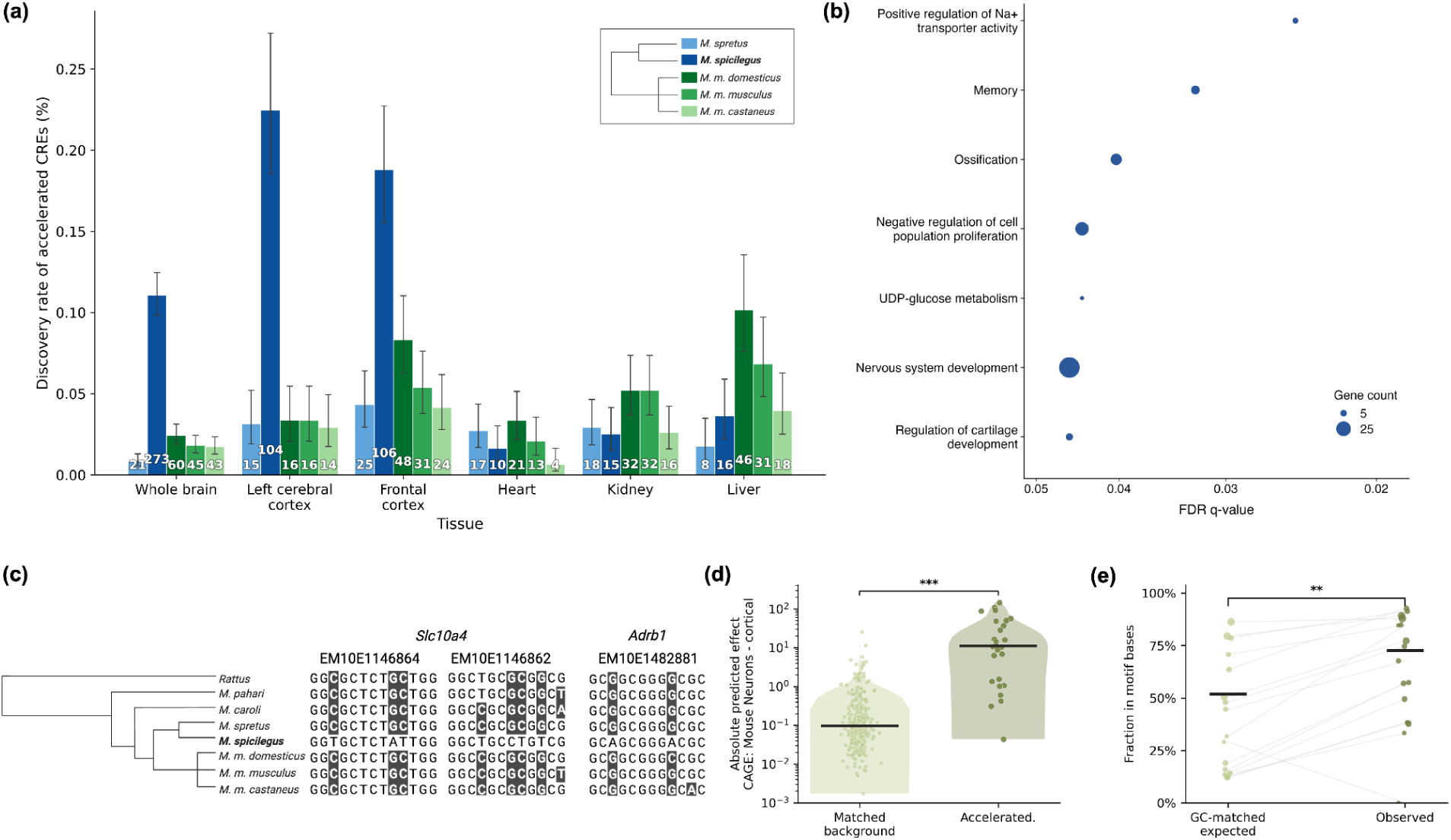
Accelerated evolution at brain-active *cis*-regulatory elements in *M. spicilegus*. Shown are results from tests of *Mus* genomes with the phyloP method for lineage-specific acceleration in candidate *cis*-regulatory elements (cCREs). (a) Each group of bars reports results of analyses of cCREs defined from assays of DNAse hypersensitivity in the indicated tissue from lab mice. In a given group, each bar reports the percentage of cCREs whose sequence exhibited significant acceleration on the indicated *Mus* lineage. Counts above bars indicate the number of cCRE hits. Error bars show Wilson 95% binomial confidence intervals. (b) Each row reports a Gene Ontology (GO) Biological Process term whose members were enriched for evidence of accelerated evolution on the *M. spicilegus* lineage at cCREs defined based on DNAse hypersensitivity in whole-brain samples (whole-brain cCREs). The *x*-axis reports -log10 Benjamini-Hochberg q-value, with larger values indicating stronger enrichment. Dot size represents the number of overlapping genes in each GO term. Shown are a subset of terms after filtering out broad, redundant parent GO terms; see Supplementary Table 4 for complete output. (c) Examples of loci with inference of accelerated evolution in *M. spicilegus* at whole-brain cCREs. *Slc10a4* is involved in motivation, reward, arousal and exploratory behavior; *Adrb1* mediates heart rate and sleep-wake regulation. (d) The *y*-axis value of each point reports the prediction, from the Enformer package, of the impact on expression in cortical neurons of *M. spicilegus*-unique alleles in a whole-brain cCRE. Columns report sets of cCREs with evidence of accelerated evolution on the *M. spicilegus* lineage (Accelerated) and a null control (Matched background). Black bars mark medians. (e) The *y*-axis value of each point reports the percentage of nucleotides in a whole-brain cCRE that overlap with at least one predicted transcription factor binding motif. Each pair of connected points reports results for two groups of cCREs with similar GC content: a set with evidence of accelerated evolution on the *M. spicilegus* lineage (Accelerated) and a GC-controlled null. Black bars mark medians. **, *p* < 0.01. ***, *p* < 0.001. Alt text: Five-panel figure characterizing *M. spicilegus* accelerated *cis-*regulatory elements. Across tissues, *M. spicilegus* shows the highest discovery rate of accelerated regulatory elements in the whole brain, cerebral cortex, and frontal cortex, with much smaller differences in heart, kidney, and liver. Accelerated brain elements are associated with gene ontology terms including nervous system development and memory. Representative sequence alignments show *M. spicilegus* specific substitutions near *Slc10a4* and *Adrb1*. Enformer predictions show larger regulatory effects for accelerated elements than matched background elements, and accelerated elements contain more motif bases than expected from GC-matched comparisons.

Inspection of the brain-active cCREs subject to accelerated evolution in *M. spicilegus* revealed many upstream of genes encoding regulators of neuronal differentiation, growth and maturation (*Sox5, Nsd1, Srgap1, Foxg1*) (Hettige and Ernst 2019; Lucas and Hardin 2017; Medina-Menéndez et al. 2025; Hamagami et al. 2023) as well as cellular and synaptic function and neural plasticity (*Slc10a4, Syt11, Rims1, Nectin3, Cdkl5, Tacr1, Nrxn2*) (Haile et al. 2023; Garcia-Recio and Gascón 2015; Zhu and Xiong 2019; Wu et al. 2022; Lonart 2002; C. Wang et al. 2018; Melief et al. 2016) (Figure 2c and Supplementary Table 3a). Some were particularly exciting owing to the evidence for their function in regulating cortical plasticity and excitation-inhibition balance (*Npas4*) (Coutellier et al. 2012) or the evolution of human neoteny (*Dock3, Srgap1*) (Fu et al. 2020; Yam et al. 2026; Namekata et al. 2010; Lucas and Hardin 2017). Likewise, unbiased analysis revealed enrichment of annotated functions relevant to behavior and development among genes whose brain-active cCREs had evolved uniquely in *M. spicilegus* (Figure 2b). The latter included the Gene Ontology term GO:0007613, which had the assigned name Memory but encompassed a diverse range of neuronal biology (Supplementary Table 4). Together, these results highlighted embryonic development of neurons and their maturation, as well as neuronal cell biology more broadly, as prominent targets of *cis-*regulatory evolution in the *M. spicilegus* lineage. A handful of genes with endocrine function also exhibited accelerated *cis*-regulatory divergence in *M. spicilegus* (*Hsd17b2, Pgrmc1, Pttg1ip, Adrb1*) (Böck et al. 2025; Read et al. 2014; Bashour and Wray 2012; Zhongyi et al. 2007) (Supplementary Table 3a).

### Pervasive *cis*-regulatory tuning by *M. spicilegus* in the cerebral cortex

We next sought neuroanatomical clues to the potential impact of *cis*-regulatory sequence divergence in *M. spicilegus*. Toward this end, we applied our method for unbiased analysis of cell-type-specific expression in the Allen Brain Atlas to the genes with accelerated evolution in brain-active cCREs. This approach detected enrichment for peak expression by our focal genes in the prelimbic area of the medial prefrontal cortex, the anterior cingulate area, and several other regions of the cerebral cortex associated with cognition, driven in part by signal from neural maturation and plasticity factors (Supplementary Figure 1b and Supplementary Table 2). This represented a contrast with the trend for expression peaks in the hypothalamus that we had noted among genes with derived amino-acid variants in *M. spicilegus* (Supplementary Figure 1a). We conclude that *cis*-regulatory variants by *M. spicilegus* may be heavily represented in cerebral cortex genes and thus are of particular interest as candidate determinants of changes in adolescent cortical maturation and the mechanisms and regulation of mound-building.

We hypothesized that the derived alleles we had detected in brain-active cCREs in *M. spicilegus* were likely to act in part by modulating mRNA expression. As a first test of this notion, we harnessed the Enformer package (Avsec et al. 2021), using models of sequence-to-expression relationships from distinct mouse cell types to predict the transcriptional effect of replacing the ancestral *Mus* allele with that from *M. spicilegus* at each brain-active cCRE in turn. Several cell-type models emerged as consistently predicting robust allele replacement effects in brain-active cCREs subject to accelerated evolution in *M. spicilegus,* but not in matched brain-active cCRE controls which accounted for substitution spectra (Supplementary Table 6 and Methods). The strongest such effects of *M. spicilegus* variants were predicted by Enformer models trained on cortical neurons from lab mice (Figure 2d and Supplementary Table 6). The mechanism likely involved transcription-factor binding, as binding sites predicted via classical bioinformatics were more likely to be perturbed by *M. spicilegus* alleles at our focal cCREs than those in matched controls (Figure 2e and Supplementary Figure 3). These results provided further support for the inference that evolution had preferentially tuned brain-active cCREs that impact neuronal gene expression in *M. spicilegus*, especially in the cerebral cortex.

The above analyses harnessed cCREs defined from chromatin profiling of whole-brain samples in two-month-old lab mice. Given the clues they had yielded to evolution of gene regulation in the cerebral cortex in *M. spicilegus,* we next focused on cortex-active loci more directly. Our approach was to revisit the search for sequence-based evidence for accelerated evolution, this time via analysis of a set of 46,309 and 56,387 cCREs defined from DNAse hypersensitivity assayed specifically in the left and frontal cortex respectively, in two-month-old lab mice (Moore et al. 2026). Applying our phyloP test for non-neutral variation to sequences from these cortex-active cCREs, we detected 178 elements with evidence for accelerated evolution in *M. spicilegus* (Figure 2a and Supplementary Table 5). Of these, almost all had originally emerged as hits in our analysis of whole-brain cCREs (Supplementary Table 3a and Supplementary Table 5), indicating that the latter signal was dominated by elements active in the cortex. We conclude that evolution has retooled *cis*-regulatory machinery in *M. spicilegus* especially avidly in the cortex, and speculate that these changes may play a causal role in the species unusual photoperiod sensitive adolescent development and/or mound-building behaviors.

Given that mound-building and related behaviors, as well as physiological phenotypes, distinguishing *M. spicilegus* from its relatives manifest at times that span the adolescent transition (Cryns et al. 2022; Brust et al. 2015), we hypothesized that they could have arisen in part via changes to gene regulation active at this point in development. To explore this, we further extended our analyses of cortex-active cCRE sequences to those detected in a time course of chromatin assays across development of lab mice (Moore et al. 2026). The results revealed the strongest evidence for accelerated evolution in *M. spicilegus* among loci active in cerebral and frontal cortices at 25 days through 2 months (the adolescent period in lab mice) and the weakest signal in cCREs active right after birth (the neonatal and juvenile period) (Figure 3a-c). No such developmental signal emerged from sequence analyses of cCREs defined across time courses of activity in the cerebellum and hippocampus, though we again detected more evidence for divergence in *M. spicilegus* than in other species at these loci (Figure 3d-e and Supplementary Table 5).

**Figure 3.**
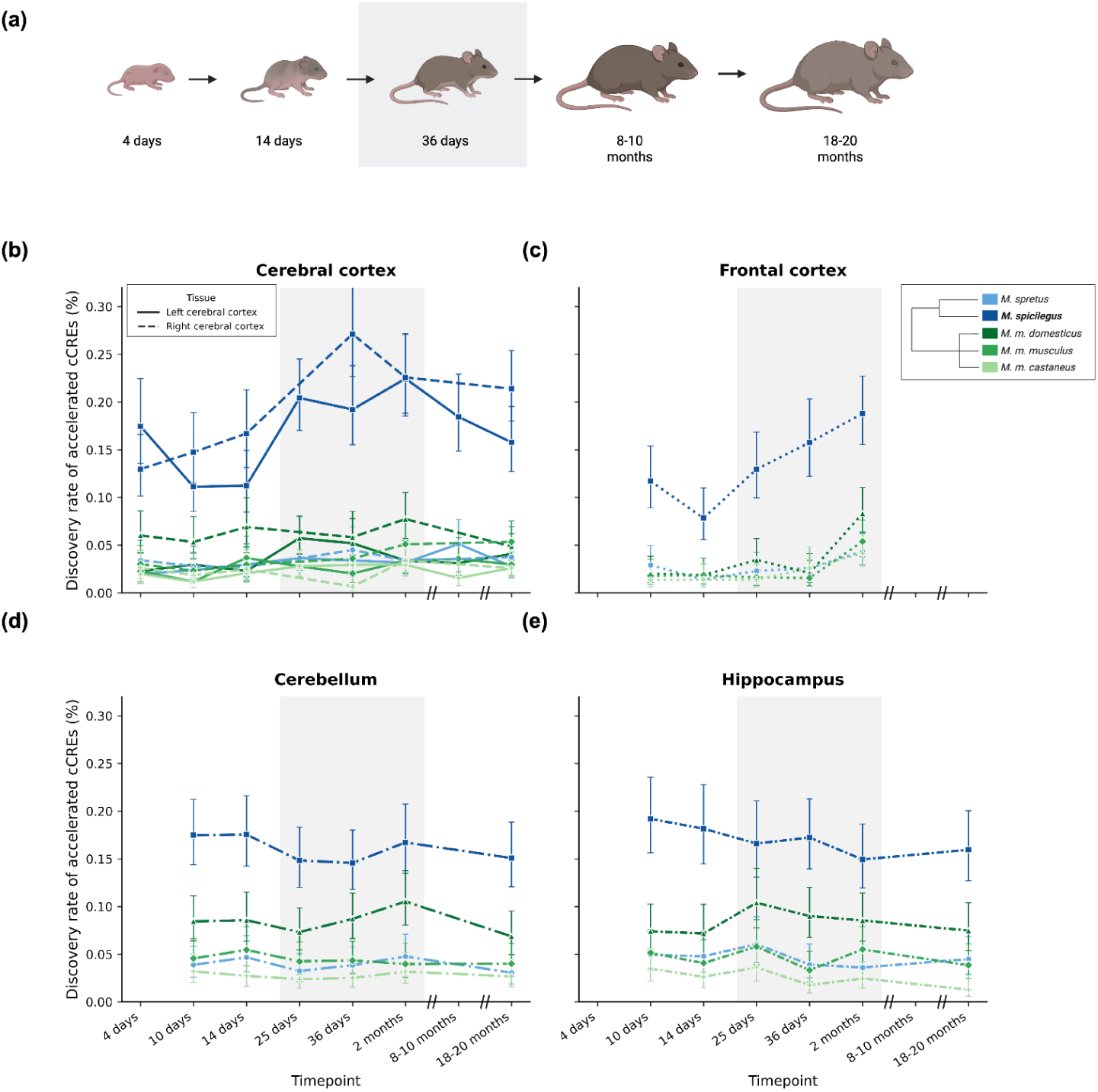
Accelerated evolution in *M. spicilegus* at *cis*-regulatory elements active in brain regions during the juvenile-adolescent-young-adult transition. (a) Stages of mouse development; grey marks the adolescent period, a transitional phase between the juvenile and young adult stages which is observed in both mice and humans (Lin and Wilbrecht 2022). Created in BioRender. Baylis, M. C. (2026) https://BioRender.com/ag6m33t. In (b)-(e), a given panel reports results from tests of *Mus* genomes with the phyloP method for lineage-specific acceleration, at cCREs defined from assays of DNAse hypersensitivity in the indicated tissue from lab mice at the age on the *x*. The *y*-axis reports the percentage of cCREs whose sequence exhibits significant acceleration on the indicated *Mus* lineage. Error bars show Wilson 95% binomial confidence intervals. Across the grouped juvenile-to-young-adult window in the cerebral cortex and the frontal cortex, discovery rates of accelerated evolution at cCREs were elevated in *M. spicilegus* relative to other *Mus* (Fisher’s exact *p* = 9.8 × 10^−4^). In a given panel, shown are results from a subset of loci after filtering out those identified as cCREs at more than 80% of timepoints, which were unlikely to have a development-specific function; see Supplementary Table 5 for complete output. Alt text: Five panel figure showing developmental patterns of accelerated *cis-*regulatory elements across mouse brain regions. A schematic depicts sampled ages from 4 days through 18-20 months. Line plots compare discovery rates across *Mus* lineages and developmental time points in cerebral cortex, frontal cortex, cerebellum, and hippocampus. *M. spicilegus* consistently shows higher acceleration rates than other lineages, with particularly elevated rates in cerebral and frontal cortex during postnatal and adult stages.

Thus, the *cis*-regulatory elements changing in *M. spicilegus* in the cerebral cortex tend to be those that are active from adolescence onward—providing further support to a model in which these loci govern adolescent mound-building or the seasonal regulation of the pace of adolescent development, dispersal and reproduction in this species.

### Divergent brain expression in *M. spicilegus*

As a complement to our sequence analyses, we expected that direct analysis of species variation in mRNA expression could reveal additional clues to the molecular mechanisms of trait divergence in *M. spicilegus*. To explore this, we harnessed RNA-seq profiles from whole-brain sampling of animals of five *Mus* species reared to 10 weeks in consistent conditions and a 12:12 photoperiod (Zhang et al. 2024). We subjected each gene’s worth of data to tests for lineage-specific expression divergence that exceeded neutral phylogenetic expectations under a Brownian-motion model (Felsenstein 1985; Zhang et al. 2024). Conforming to our prediction, we detected more genes with significant, non-neutral expression divergence in the brain in *M. spicilegus* than in any other species, a total of 1500 such hits (Figure 4a and Supplementary Table 8). No such excess in signal for lineage-specific expression in *M. spicilegus* was apparent from analyses of heart, kidney, or liver transcriptomes (Supplementary Figure 4), paralleling the trends we had seen in our tests of sequence divergence (Figure 2). For neuroanatomical insights beyond the whole-brain level, we applied our paradigm for analysis of cell-type-specific expression in the Allen Brain Atlas to the hits from our screen for lineage-specific programs in the brain in *M. spicilegus*. The results revealed that the latter were enriched for expression in the isocortex, the anterior cingulate area, and other domains of the cerebral cortex, attributable in part to genes implicated in neural development and synaptic plasticity (Supplementary Figure 1 and Supplementary Table 2). This trend is consonant with our findings from analysis of *cis*-regulatory sequence (Figures 2-3), further supporting our model of the cerebral cortex genes as a hotspot for expression programs unique to *M. spicilegus*.

**Figure 4.**
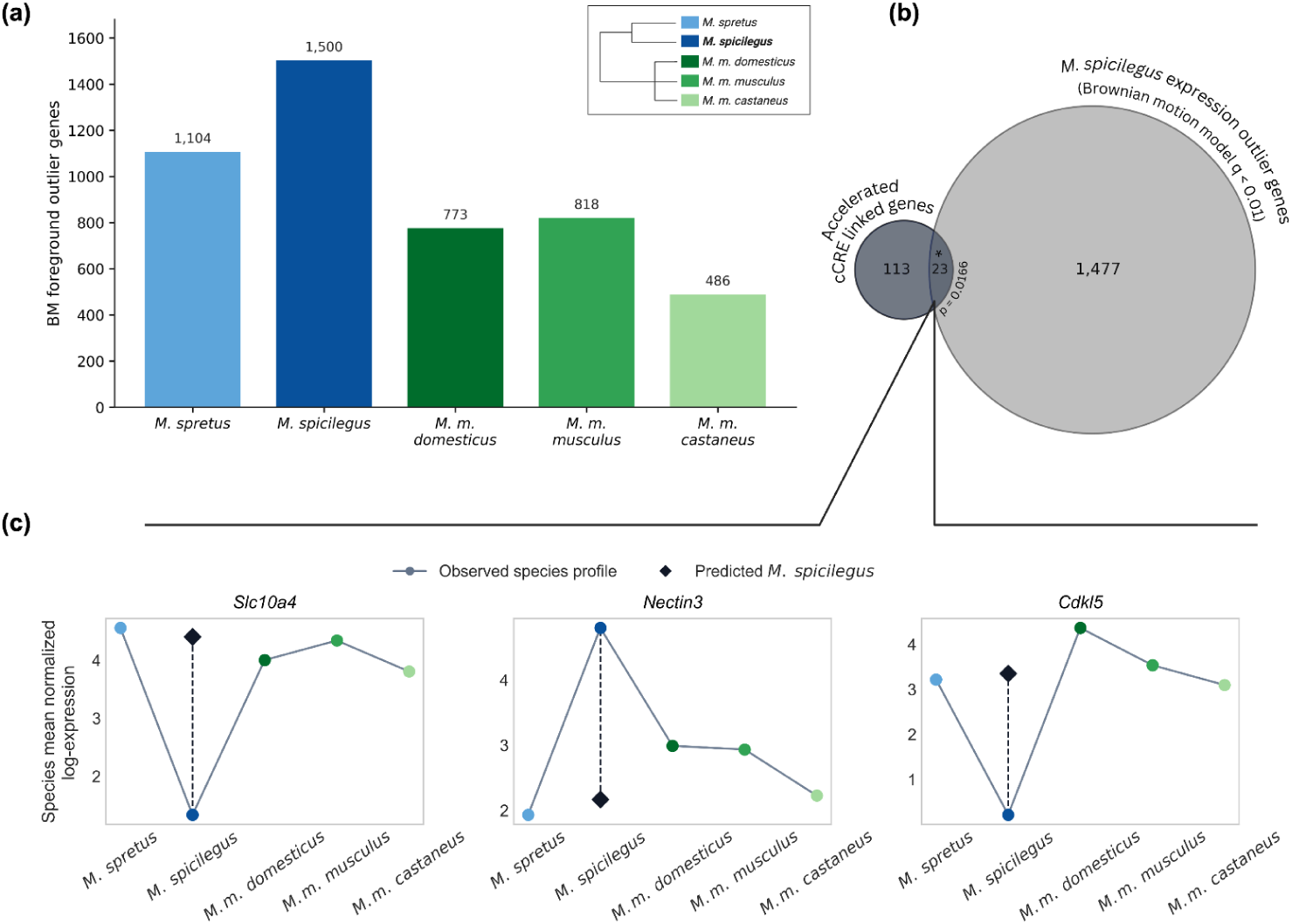
Accelerated divergence in brain expression in *M. spicilegus.* Shown are results from tests for genes exhibiting expression divergence in a given mouse species more dramatic than that expected under a Brownian-motion model, measured from whole-brain samples at age 60 days. (a) Each bar reports the number of genes with significant brain expression divergence in the indicated species. (b) Circles report numbers of genes associated with whole-brain cCREs exhibiting accelerated evolution in *M. spicilegus* (dark) and genes exhibiting divergent brain expression in *M. spicilegus (*light); intersection between the sets is significant at one-sided Fisher’s exact *p* = 0.0166. (c) Each panel reports results for an example gene whose whole-brain cCRE sequence and brain expression both exhibited accelerated evolution in *M. spicilegus.* In a given panel, each circle reports whole-brain expression from the indicated species, and the diamond reports the expression level predicted for *M. spicilegus* under the Brownian-motion null. See Supplementary Figure 5 for the complete set of overlapping genes from (b). Alt text: Three panel figure comparing evolutionary shifts in brain gene expression among *Mus* lineages. *M. spicilegus* has the largest number of brain expression outlier genes. A Venn diagram shows limited overlap between *M. spicilegus* expression outliers and genes associated with accelerated *cis-*regulatory elements. Species level expression plots for *Slc10a4*, *Nectin3*, and *Cdkl5* illustrate observed *M. spicilegus* expression relative to values predicted from the other species.

We next returned to the question of the impact of *cis*-regulatory alleles in *M. spicilegus* that had emerged from our sequence-based tests (Figures 2-3). To integrate these loci with direct measurements of mRNA expression, we compared the genes top-scoring in our tests for divergence in *M. spicilegus* from our two approaches: hits from phyloP applied to whole-brain cCREs on the one hand and from Brownian-motion analysis of transcriptomes on the other. The results revealed overlap at a level that was small in magnitude but robustly significant (Figure 4b-c and Supplementary Table 8), establishing the respective cCREs as particularly well-supported candidate drivers of expression change in *M. spicilegus*. These cases included *Cdkl5, Maged1*, *Nectin3,* and *Slc10a4,* genes impacted in neural development, neurotransmission, dendritic spine density, and synaptic plasticity (Figure 4c, Supplementary Figure 5, and Supplementary Table 8) (J. Yang et al. 2015; Della Sala et al. 2016; X.-D. Wang et al. 2013; Larhammar et al. 2015; Zhu and Xiong 2019; J. Wang et al. 2023; Tomorsky et al. 2020; Wu et al. 2022). Of the remainder, genes with divergent expression in *M. spicilegus* but no detectable non-neutral sequence change in our cCRE set would be expected if they were controlled by *trans*-acting regulatory variation instead. Variant cCREs at genes with no detectable expression divergence in the whole brain could be expected under a model of tissue-specific or post-transcriptional effects. That said, the most salient conclusion from our analyses was that *M. spicilegus* expresses unique programs in the brain not seen in other *Mus*, and that variant brain-active cCREs are likely players in their genetic architecture.

### *M. spicilegus* candidate genes overlap gene programs associated with independently evolved complex behavior

For further insights into the potential phenotypic impact of molecular changes in *M. spicilegus*, we now sought to interpret them with respect to roles of the respective genes in the brain in other animal systems. We considered a set of genes whose protein products were detected in the post-synaptic density of rodent and human excitatory neurons in the neocortex; a set of genes at which copy number variation is associated with autism risk in humans; and expression correlates of bower-building behavior in cichlid fish (Figure 5 and Supplementary Table 9) (Olson et al. 2015; Johnson et al. 2023; SPARK 2025; L. Wang et al. 2023). We assessed each cohort in turn for overlap with genes whose brain-active cCREs had signal for accelerated sequence evolution in *M. spicilegus* and, separately, with genes exhibiting lineage-specific expression in *M. spicilegus*. Results revealed overlap in almost every comparison (Figure 5 and Supplementary Table 9). Of particular interest was the Rho family signaling pathway active in the post-synaptic density (including *Dock3* and *Srgap1*), which mediates the delayed maturation of human neurons in the neocortex relative to other species (Olson et al. 2015; Johnson et al. 2023; SPARK 2025; L. Wang et al. 2023) and could well carry out a similar role in *M. spicilegus* animals born in autumn that show protracted development over the winter. Writ large, our results make clear that genes subject to regulatory divergence in *M. spicilegus* are associated with neuronal development, cognition, and motivated behavior across animal models, further supporting the inference of their roles in related traits in *M. spicilegus*.

**Figure 5.**
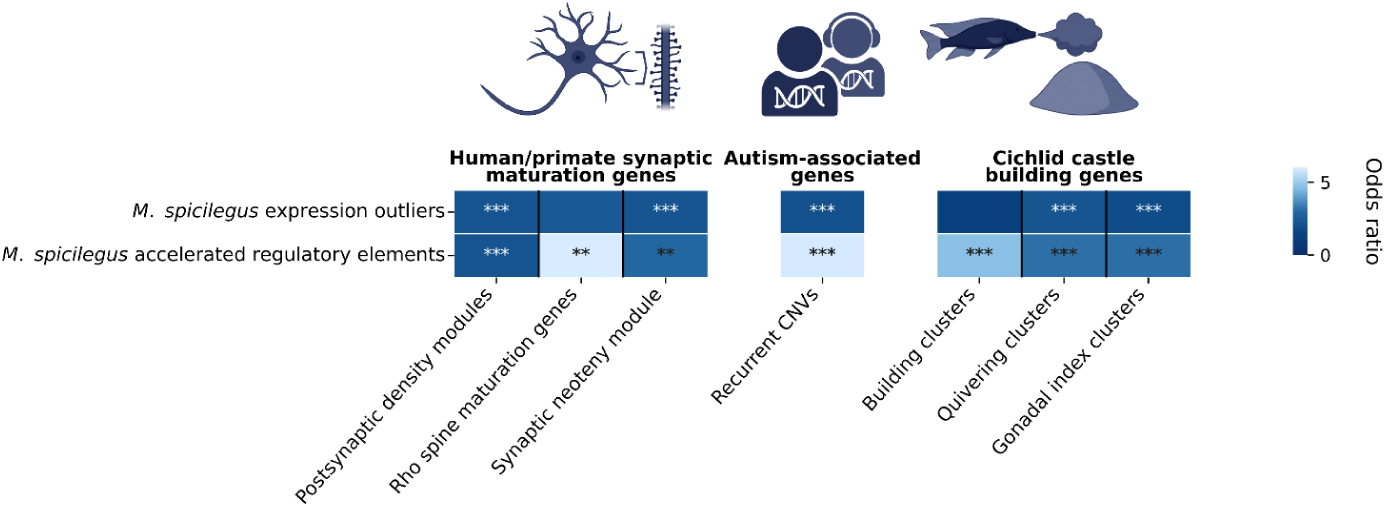
Accelerated sequence and expression divergence in *M. spicilegus* at genes associated with neurodevelopment and behavior. Each heatmap denoted by a distinct cartoon reports the degree of overlap between two gene sets: genes with evidence for divergence in *M. spicilegus* (in expression, top row, or whole-brain cCRE sequence, bottom row) and genes implicated in neurodevelopment or behavior in another organismal system. Each cell reports the odds ratio (color) and significance (asterisks) of the intersection of the gene sets in a Fisher’s exact test. In the first heatmap, columns report analysis of, from left to right, genes encoding proteins detected in the post-synaptic density of human cortical neurons; a subset of the latter annotated as RhoGEFs and RhoGAPs; and a separately defined subset whose protein levels were higher in post-synaptic density from human cortical neurons than those from macaque or mouse, from (L. Wang et al. 2023). In the second heatmap, the column reports analysis of genes at which copy-number variation is associated with autism in human patients; from (SPARK 2025). In the third heatmap, columns report analysis of, from left to right, genes whose expression is associated with bower-building behavior, fin-quivering behavior, and gonadal size in courting male cichlid (*Mchenga conophoros*) fish, from (Johnson et al. 2023). **, *p* < 0.01. ***, *p* < 0.001. Alt text: Heat maps showing enrichment of *M. spicilegus* brain expression outliers and accelerated regulatory elements in previously defined gene sets. Enrichment is shown for human and primate synaptic maturation gene modules, autism-associated genes, and cichlid castle-building gene clusters. Multiple significant enrichments occur across both *M. spicilegus* expression outliers and accelerated *cis-*regulatory elements.

## Discussion

Against the backdrop of successful quantitative-genetic studies of behavior within mammalian populations, dissecting the genetics of behavioral variation between species remains an ongoing challenge (Roux et al. 2014; Weber et al. 2013; Hu and Hoekstra 2017; Hu et al. 2022; York et al. 2018; Johnson et al. 2023; Kapheim et al. 2015; Doan et al. 2016; Capra et al. 2013; Pollard et al. 2006). The *Mus* genus is a powerful model for interspecies genetics, in which resources from lab mice can be brought to bear in the study of trait variation from the wild (Larson et al. 2018; Keithley et al. 2004; Logan et al. 2013; Song et al. 2011). In this work, we have pioneered genomic analyses in *M. spicilegus* to begin to shed light on the evolutionary forces and molecular mechanisms underlying its unique phenotypes. In analyses of amino-acid variation, brain-active cCRE sequence, and brain mRNA expression, we have shown that non-neutral divergence in *M. spicilegus* far outstrips that in its sister species, strongly suggestive of a history of adaptation in hypothalamic and cortical functions. And the number of divergent loci (>200 in the case of sequence variation, and almost an order of magnitude more with derived expression) points to a complex genetic basis for *M. spicilegus*’ traits.

### Seasonality of sexual maturation and the adolescent transition

In the search to understand *M. spicilegus’* evolution, reproductive physiology represents a key piece of the puzzle, given the overwintering in mounds and long delay to dispersal and of reproduction seen in autumn-born *M. spicilegus* but not other *Mus* (Cryns et al. 2022; Lafaille et al. 2015; Poteaux et al. 2008; Gouat et al. 2003b). In this capacity for protracted reproductive development, *M. spicilegus* resemble Siberian hamsters, which delay puberty over the winter to breed in the spring (Paul et al. 2018). The specific signals and mechanisms that arrest reproductive development in *M. spicilegus* in the autumn remain unknown. Our results have revealed prime candidate determinants at the molecular level, especially the amino-acid divergence in *M. spicilegus* in genes implicated by other studies in pubertal timing (*Tmem108*) (Rezende et al. 2023), specific aspects of male sex behavior (*Sytl4*) (X. Xu et al. 2012), and major pathways implicated in circadian function, development, and even aging (*Per3, Sdc3, Vglut2*, *Zmpste24*) (Archer et al. 2018; Strader et al. 2004; Worman and Michaelis 2023; Nakamura et al. 2005; Arokiasamy et al. 2019; Matsumura and Akashi 2017). Furthermore, the patterns of variation in these genes raise the intriguing possibility that endocrine functions of the hypothalamus in *M. spicilegus* could have been tuned by evolution preferentially via protein-coding changes, with *cis*-regulatory tuning as a more minor contributor. If so, it would echo classic discoveries of protein variation in circadian clock factors that underlies naturally varying behaviors in fruit flies and humans (Wheeler et al. 1991; Y. Xu et al. 2005). Whether coding or non-coding, differences in hypothalamic function are a strong candidate for determinants of seasonal timing of sexual reproduction in *M. spicilegus*, based on the current understanding of seasonal breeding in other animals (Stevenson et al. 2026; Nishiwaki-Ohkawa and Yoshimura 2016; Clarke et al. 2009; Weems et al. 2015; Prendergast et al. 2000; 2002).

Interestingly, however, endocrine genes did not feature among the most pervasive patterns we noted in regulatory divergence of *M. spicilegus*. Rather, the latter signal was dominated by genes controlling neuronal development and plasticity, particularly in the cerebral cortices with dynamic regulation in the cortex generally and the frontal cortex specifically during adolescence. Most notable were cases of *cis*-regulatory change in *M. spicilegus* at drivers of axon and dendrite growth, synapse formation, activity-dependent transcription, and excitatory neurotransmission and excitatory/inhibitory balance. We also observed variants in genes controlling the brain’s specialized neuromodulatory neurons—including factors that act in the function of their axon terminals as well as neuromodulatory neurotransmitter receptors. Under one compelling model, some of these changes could act in part to enable autumn-born *M. spicilegus* to delay maturation of some cortical circuits and retain a state of relative neoteny through the winter (Piekarski et al. 2017; Cryns et al. 2022; Bhatt et al. 2009). Such a program could mirror the delayed, neotenic synaptic maturation in humans relative to other mammals (Charrier et al. 2012; L. Wang et al. 2023). This idea is supported by our finding of significant overlap between genes whose regulation has diverged in *M. spicilegus* and genes implicated in human cortical synapse maturation (*e.g. Dock3*). Likewise, the particularly strong sequence divergence that we have noted in *M. spicilegus* at cortical regulatory elements active during adolescent development, relative to those active earlier and later in development, can be interpreted as further support for a potential role for these variants in regulation of the pace of adolescence or biological milestones within the adolescent transition.

### Mound-building and dispersal

Even if cortical maturation is delayed in *M. spicilegus* in the autumn, other cognitive functions must be preferentially active in this season to enable social mound-building and overwintering in mounds. This suite of behaviors makes autumn-born behavioral phenotypes of *M. spicilegus* quite different from those of spring-born animals of the same species as well as from other *Mus* (Hurtado et al. 2013; Szenczi et al. 2011; Poteaux et al. 2008; Gouat et al. 2003b; Garza et al. 1997). It is tempting to speculate that selective regulation of the neural development and plasticity genes at which we have noted variation in *M. spicilegus* may contribute to sociality and/or building in autumn-born members of the species; primary candidates for such a role include *Npas4 (*Coutellier et al. 2012*), Nectin3* (Tomorsky et al. 2020; Wu et al. 2022; X.-D. Wang et al. 2013; Ramamoorthi et al. 2011; Weng et al. 2018; X. Li et al. 2014; Melief et al. 2016), *Cdkl5* (I.-T. J. Wang et al. 2012), and *Nrxn2* (Haile et al. 2023). Consonant with these observations in the *M. spicilegus* genome, studies of *Peromyscus* deer mice (Hu et al. 2022) and cichlid fishes (York et al. 2018; Johnson et al. 2023) suggest that both have used *cis*-regulatory change in neurodevelopmental loci and circuit-level regulators to encode both social and building phenotypes. And the broader literature has implicated expression divergence in the evolution of many other cases of cognition, most notably sociality in insects, sociality in voles, and higher reasoning in humans (Jernigan et al. 2023; Phelps et al. 2017; Kapheim et al. 2015; Doan et al. 2016; Capra et al. 2013; Pollard et al. 2006).

Additionally, given that in *M. spicilegus* the age of dispersal depends on the season of birth, the developmental milestone of dispersal is likely to be dynamically regulated, representing a powerful model for study of the neural basis of dispersal behavior (Groó et al. 2018; Lin and Wilbrecht 2022; Cryns et al. 2022). Dopamine and other neuromodulators play a major role in shaping motivated behavior and therefore are likely important in regulating the arousal and drive necessary for dispersal. As such, we can speculate that the molecular changes we have identified in *M. spicilegus* in genes that impact dopamine neuron biology, such as *Syt11* (C. Wang et al. 2018), *Slc10a4* (Melief et al. 2016), *Maged1* (J. Wang et al. 2023), or *Adrb1* (Böck et al. 2025), a major adrenergic receptor, may be differentially regulated by season of birth in this species to mediate dispersal. In sum, we propose that the molecular changes we have observed in *M. spicilegus* can be harnessed as clues to the neural mechanisms that regulate dispersal and other motivated behaviors central to adolescent development, to guide future study of the impact of rearing photoperiod on *M. spicilegus* neurobiology.

### Conclusions

The trends traced by our analyses of the *M. spicilegus* genome and transcriptome provide key insights into the history of natural selection on CNS function as a force underlying the divergence of this species. And the cases of molecular variation that we highlight represent candidate determinants of unique traits in *M. spicilegus*, to be validated by future advances in the field. We expect that *M. spicilegus* will continue to be a powerful model for studies in mammalian evolution, in which resources from laboratory mice can be marshaled to understand how nature builds new facets of behavior and development in the wild.

## Methods

### 1. Species, phylogeny, and ortholog selection

Orthologous coding sequences were compiled across eight Murinae taxa. Multiple sequence alignments were derived from the 21-way Enredo-Pecan-Ortheus (EPO) alignments corresponding to Ensembl Compara (release 112) (Cunningham et al. 2022; Paten, Herrero, Fitzgerald, et al. 2008; Paten, Herrero, Beal, et al. 2008). For our analyses, we restricted the phylogeny to eight focal taxa representing one outgroup (*Rattus norvegicus*), three wild non-musculus *Mus* species (*M. pahari, M. caroli, M. spretus*)*, M. spicilegus,* and three representative *M. musculus* subspecies strains (*M. m. domesticus WSB/EiJ, M. m. musculus PWK/PhJ, M. m. castaneus CAST/EiJ*). Alignments were downloaded in EMF format and converted to MAF using the Ensembl Compara emf2maf parser (Cunningham et al. 2022; Paten, Herrero, Fitzgerald, et al. 2008; Paten, Herrero, Beal, et al. 2008). Alignments were grouped by mouse reference chromosome. Coding-sequence analyses used GRCm39 coordinates, whereas cCRE analyses used mm10 coordinates to match the ENCODE SCREEN mouse cCRE annotations. For cCRE-based phylogenetic analyses, mm10 cCRE intervals were used as the reference coordinate anchors for extracting orthologous alignment blocks from the corresponding mouse-referenced EPO alignments.

### 2. Multiple sequence alignment quality control

Because lineage-specific acceleration can be inflated by assembly, orthology, or local alignment artifacts, coding and *cis*-regulatory alignments were filtered before statistical analysis and candidate interpretation (Fletcher and Yang 2010; Jordan and Goldman 2012; Pollard et al. 2010). Elements were required to have sufficient taxon representation, an identifiable foreground sequence, equal alignment lengths across retained species, and callable foreground bases. Columns with gaps or ambiguous bases in the focal foreground were excluded from foreground private substitution summaries.

In tabulating alleles unique to *M. spicilegus*, for a given alignment we calculated foreground callable coverage, foreground private substitution count and rate, the maximum number of foreground private substitutions within any 10 bp window, the longest local serial cluster of private substitutions separated by no more than 2 bp, and the number of foreground private gaps. A foreground private substitution was counted only when the foreground base was callable, at least five nonforeground taxa had callable bases, the nonforeground taxa supported a single majority base at a frequency at least 0.75, and the foreground base differed from that majority state. Candidate alignments were eliminated from analysis if they showed strong error-like patterns, namely private substitution rate greater than 0.08, at least 25 foreground private substitutions, at least seven private substitutions in any 10-bp window, or at least three foreground private gaps.

### 3. Coding sequence tests for selection (PAML)

Protein-coding evolution was tested with codon models in PAML codeml v4.10.7 (Z. Yang 1997; 2007). For a given gene, coding sequences of each ortholog were translated (MAFFT v7.526), aligned at the amino-acid level, translated to codon alignments to preserve the reading frame (PAL2NAL v14), and filtered to remove frame-disrupted, stop-containing, or poorly-aligned sequences (Tree topology was as in Figure 1a) (Katoh and Standley 2013; Suyama et al. 2006).

For each gene, each focal *Mus* lineage was labeled as the foreground branch in turn. We implemented codeml allowing the nonsynonymous to synonymous rate ratio (ω = dN/dS) to vary on the foreground branch relative to the rest of the tree. For each gene, we compared the alternative model, a two-ratio branch model allowing ω foreground to differ from ω background with ω foreground estimated freely (including values > 1), with the one ratio null model in which a single ω was estimated across all branches. Likelihood-ratio statistics were calculated as twice the log likelihood difference between alternative and null models, and p-values were adjusted within each foreground using the Benjamini-Hochberg false discovery rate procedure (Benjamini and Hochberg 1995).

### 4. cCRE definition and gene annotation

Our catalogs of candidate *cis*-regulatory elements (cCREs) from lab mice were from the ENCODE Registry (the SCREEN database; Ensembl/ENCODE release aligned to mm10) (ENCODE Project Consortium et al. 2020). We retained for analysis all loci called as active by the ENCODE Registry based on tissue-specific DNase-seq signal (within-biosample Z-score > 1.64). For gene assignments, we used ENCODE cCRE-to-gene linkage annotations. cCRE coordinates were obtained in mm10 reference coordinates and used as anchors for orthologous alignment extraction (see section 2).

cCREs active in whole-brain, heart, liver, and kidney were from SCREEN calls from DNAse hypersensitivity measured at P60. Left cerebral cortex-active cCREs were from DNAse hypersensitivity measured at postnatal day 4, day 10, day 14, day 25, day 36, 2 months, 8-10 months, and 18-20 months. Right cerebral cortex-active cCREs were from DNAse hypersensitivity measured at day 4, day 10, day 14, day 36, 2 months, and 18-20 months. Cerebellum and hippocampus-active cCREs were from DNAse hypersensitivity measured at day 10, day 14, day 25, day 36, 2 months, and 18-20 months. Frontal cortex-active cCREs were from DNAse hypersensitivity measured at day 10, day 14, day 25, day 36, and 2 months. For a given time course in Figure 3b-e, cCREs active in more than 80% of developmental timepoints were classified as unlikely to have developmental-specific functions and excluded from analysis.

To design GC-aware control procedures, for each cCRE alignment we counted foreground private sites and calculated the fraction represented by GC changes. cCREs with at least five GC private sites and with GC changes comprising at least 50% of all strict private foreground sites were excluded from further analyses. For the remainder, we stratified substitution sites into bins on the basis of base-change class, local GC content around the mutated base, and whole CRE GC content. We used these bin allocations in sections 7 and 8 below.

### 5. phyloP subtree tests for regulatory acceleration

We used phyloP subtree tests to identify *cis*-regulatory elements with significantly accelerated sequence evolution on specific foreground lineages, indicating candidate regulatory regions that may have been evolutionarily modified in those lineages (Pollard et al. 2010; Cooper et al. 2005; Siepel et al. 2006). The phylogenetic tree used for phyloFit and phyloP analyses corresponded to the Ensembl EPO species tree with branch lengths estimated under the EPO framework. The topology reflects the expected Murinae relationships, with *M. spicilegus* sister to *M. spretus*, nested within the broader *Mus* clade, and *Rattus norvegicus* as the outgroup. To model neutral sequence evolution, we fit a phylogenetic substitution model using phyloFit (PHAST/phyloP/phyloFit v1.5) to estimate branch lengths and substitution-rate parameters under a reversible nucleotide substitution model on the fixed species tree topology (Hubisz et al. 2011). As neutral training data, we used fourfold degenerate (4D) codon sites extracted from the same set of one-to-one orthologous genes analyzed in the coding selection scan (Chakraborty 2002). Fourfold degenerate sites were chosen because they are expected to be largely free of selective constraint at the amino-acid level and therefore approximate neutral substitution dynamics. Sites overlapping annotated regulatory regions or conserved elements were excluded from the neutral training set to reduce potential contamination by constrained sequence. The resulting neutral model captured lineage-specific background substitution rates and served as the baseline expectation for regulatory sequence evolution. Using this neutral model, we applied phyloP to test for deviations from neutrality at each cCRE alignment. phyloP computes likelihood-based statistics comparing the observed substitution pattern at a locus to the neutral expectation, producing p-values for conservation (slower than neutral evolution) or acceleration (faster than neutral evolution). We used the Siepel Pollard Haussler method, which tests whether substitution rates along a designated subtree deviate from the neutral model while conditioning on total substitution counts across the full phylogeny (Pollard et al. 2010). We used the SPH subtree test in CONACC mode (--method SPH --mode CONACC --subtree), which tests whether the substitution rate in a designated subtree differs from that expected under the neutral model, conditional on the complementary supertree. This framework allows us to identify lineage specific shifts in evolutionary rate while accounting for locus specific variation in evolutionary history across the remainder of the phylogeny.

For each cCRE, phyloP generated branch specific p-values for acceleration and conservation on the focal subtree, as well as supertree-wide statistics. p-values were adjusted across all tested elements using the Benjamini-Hochberg false discovery rate procedure (Benjamini and Hochberg 1995). Elements with FDR < 0.05 in the acceleration test for the *M. spicilegus* subtree were classified as lineage-specific accelerated cCREs.

cCREs called by this procedure as subject to accelerated evolution in *M. spicilegus* were not enriched for poor alignment, elevated gap content, or ambiguous bases, indicating that phyloP acceleration signals were unlikely to reflect local assembly or alignment artifacts. For each species, timepoint, or tissue, the accelerated CRE discovery rate was calculated as

Discovery rate (%) = 100 * significantly accelerated CREs / total CREs tested.

### 6. Gene Ontology enrichment analysis

Gene Ontology Biological Process enrichment was performed on genes linked to accelerated *M. spicilegus* whole brain cCREs using g:Profiler (gprofiler2 0.2.4 in R v4.1.1) with custom background and FDR correction (Raudvere et al. 2019). Foreground genes were defined as those linked to at least one significantly accelerated *M. spicilegus* brain cCRE, and background genes were defined as all genes linked to any brain cCRE in the same ENCODE universe. Enrichment p-values were corrected across tested GO terms using Benjamini-Hochberg FDR (Benjamini and Hochberg 1995).

### 7. Sequence-to-expression prediction with Enformer

To estimate the potential regulatory consequences of *M. spicilegus*-specific sequence variants in accelerated cCREs, we used Enformer, a deep-learning model that predicts functional genomic signals from DNA sequence (Avsec et al. 2021). We used the DeepMind Enformer model from TensorFlow Hub (https://tfhub.dev/deepmind/enformer/1), run with TensorFlow v2.21.0 and TensorFlow Hub v0.16.1 (Avsec et al. 2021). Input windows were generated from the UCSC mm10 mouse genome, and mouse output-track metadata were taken from the Basenji/Enformer targets_mouse.txt file. For each accelerated brain-active cCRE, we generated paired input sequences representing the inferred ancestral/non-*M. spicilegus* state and the *M. spicilegus*-derived state. Variant positions were defined from the multiple sequence alignment as sites at which the *M. spicilegus* base differed from the majority base among callable non-*M. spicilegus Mus* taxa. Paired sequences were otherwise identical, allowing predicted differences to be attributed to the *M. spicilegus*-specific substitutions within the cCRE. For each sequence pair, Enformer predictions were extracted for 351 mouse output tracks corresponding to neural tissues or cell contexts. Allelic effects were quantified as the difference in predicted signal between the *M. spicilegus*-derived and ancestral/non-*M. spicilegus* sequence across bins overlapping the focal cCRE. To assess whether predicted effects exceeded expectations from local sequence composition and substitution burden, accelerated cCREs were compared with matched non-accelerated brain-active cCREs selected to have similar length, GC content, and *M. spicilegus*-specific substitution count (see section 4; data are in Supplementary Figure 3 and Supplementary Table 6).

### 8. Transcription factor motif analysis

To test whether *M. spicilegus-*specific substitutions within cCREs subject to accelerated evolution altered predicted transcription factor-binding architecture, we analyzed 285 cCRE alignments with evidence for accelerated evolution in *M. spicilegus* and 285 matched background cCRE alignments with no such signal.

For each alignment, we tabulated nucleotide sites at which *M. spicilegus* had one allele and all other *Mus* had another allele, the latter representing the consensus ancestral sequence. We used this to construct a consensus ancestral sequence of the cCRE and, separately, we substituted all *M. spicilegus*-unique alleles into this consensus for a stand-alone *M. spicilegus* version of the locus. Then, we scanned each in turn in local sequence windows overlapping each difference site with JASPAR position weight matrices (Castro-Mondragon et al. 2022). Motif scores were converted to relative motif scores using the minimum and maximum possible score for each motif. A site was classified as a motif-switch site when the best-scoring motif identity differed between the *M. spicilegus* and the consensus states, and at least one state passed the relative score threshold.

To test whether *M. spicilegus* substitutions were preferentially concentrated within predicted transcription factor motif bases, we intersected substitution positions with annotated motif matches. To evaluate the significance of this overlap, because variants unique to *M. spicilegus* in cCREs with evidence for accelerated evolution showed GC-biased sequence change, we used a GC- and substitution-matched framework for permutation-based comparisons to a matched background set, as follows. Substitution sites were assigned to bins defined by local GC content surrounding the substitution and overall cCRE GC content (see section 4). Within these matched GC context strata, *M. spicilegus* labels were permuted to generate the expected number of substitutions overlapping motif bases under a GC-matched null model. Observed motif-overlapping substitutions were then compared with this null expectation. *M. spicilegus* substitutions overlapped predicted motif bases more often than expected from GC context alone (observed/expected = 1.40, permutation p = 5.0 × 10^−5^).

### 9. Allen Brain Atlas expression patterns of evolutionary candidate genes

To connect our evolutionary candidate gene sets to brain-region biology, we used the Allen Mouse Brain Atlas adult mouse *in situ* hybridization dataset to ask where these genes are expressed in the adult mouse brain. This resource provides spatially resolved expression energy measurements from 56-day-old male C57BL/6J mice, allowing us to compare expression across named brain regions and finer anatomical subregions. Expression energy reflects both the intensity and spatial extent of *in situ* hybridization signal within a structure, so higher values indicate stronger and/or broader expression in that region. Genes with no expression energy measurement in the tested region were excluded from that region set analysis.

For each gene, we compared Allen adult mouse brain ISH expression-energy values across the regions and assigned the gene’s peak to the region with the highest expression-energy value. Expression energy is a quantitative Allen Atlas summary that reflects both the intensity of hybridization signal and the spatial extent of that signal within a structure. In cases where multiple Allen experiments or section datasets were available for the same gene and region, we used the maximum expression-energy value so that each gene contributed a single value per region.

For enrichment analyses, each evolutionary candidate gene set was intersected with the Allen background matrix for the same region set. Background genes were all Allen genes with expression-energy data in that matrix after removing the candidate genes being tested. For each region, we built a 2-by-2 table comparing candidate genes peaking in that region, candidate genes peaking elsewhere, background genes peaking in that region, and background genes peaking elsewhere. One-sided Fisher exact tests were used to ask whether candidate genes were more likely than Allen background genes to peak in each region.

### 10. RNAseq processing and Brownian motion expression models

Whole-brain RNAseq data from five *Mus* taxa (*M. m. musculus*, *M. m. castaneus*, *M. m. domesticus*, *M. spicilegus*, and *M. spretus*) from mice at 10 weeks old, exposed to 12:12 photoperiod (Zhang et al. 2024), were reprocessed using a comparative expression pipeline as follows. RNAseq reads from each species were aligned to their respective genome assemblies to minimize reference mapping bias. To place expression estimates into a common gene space, annotations were standardized across species by projecting the mouse reference annotation onto each genome using Liftoff v1.6.3, which uses minimap2 v2.28 for sequence alignment (Shumate and Salzberg 2021; H. Li 2018). Gene level counts were then generated from the resulting alignments using HTSeq v2.0.3 (Anders et al. 2015). Genes were matched across species using orthologous gene symbols, and only genes successfully identified in all five taxa were retained for downstream comparative analyses. This procedure yielded a shared set of homologous genes suitable for cross-species expression comparisons.

Normalized expression values were estimated using the voom procedure in limma (v 3.48.3), which transforms count data to log scale expression values while modeling the mean-variance relationship to generate precision weights for linear modeling (Law et al. 2014; Ritchie et al. 2015). Species mean-normalized log-expression values were then calculated by averaging voom expression values across samples within each species.

To determine whether a given species exhibits gene expression more unusual than expected, given the other species and their phylogenetic relationships, we used Brownian motion modeling (Felsenstein 1985; Garamszegi 2016). For each gene, the vector of species mean expression values (*M. m. musculus, M. m. castaneus, M. m. domesticus, M. spicilegus, M. spretus*) was evaluated under a Brownian motion model using a fixed covariance matrix representing the assumed five taxon phylogeny. The conditional expectation of expression in a focal taxon was computed given the remaining taxa, and the observed minus expected residual was standardized using the conditional variance to obtain a Brownian motion residual z-score. We defined a conservative expression-outlier set using stringent filtering criteria: q < 0.01, |z| >= 3, and an absolute-Z margin >= 2 over the next-most-extreme species. These filters were chosen to prioritize genes with strong statistical support, large standardized deviations from Brownian motion expectations, and expression divergence most clearly attributable to the focal lineage rather than to comparable outlier behavior in another species.

In addition to the analytic p-values derived from the normal distribution, empirical p-values were estimated from the absolute Brownian motion residual z-scores using 200,000 draws from the absolute standard normal distribution. These empirical p-values were adjusted for multiple testing across genes using the Benjamini Hochberg procedure.

To assess the overlap between genes with evidence for accelerated evolution in *M. spicilegus* cCRE sequence and genes with expression divergence in M. spicilegus in Figure 4b, we used a one-sided Fisher’s exact test. A 2 x 2 contingency table was constructed from the number of candidate genes significant for BM extremeness, candidate genes not significant, background genes significant but not in the candidate set, and background genes neither significant nor in the candidate set. Expected overlap under the null hypothesis was calculated as the product of the candidate list background count and the size of the significant background set, divided by the total background size.

### 11. Comparison of *M. spicilegus* candidate genes with known conserved developmental programs underlying complex behavior

We tested whether *M. spicilegus* candidate genes were enriched for genes implicated in synaptic maturation, neurodevelopmental risk, and independently evolved vertebrate behavioral plasticity. In one analysis, candidates were genes identified as lineage-specific expression outliers in *M. spicilegus* from Brownian-motion models; in a separate analysis, candidates were linked to candidate *cis*-regulatory elements with accelerated evolution in *M. spicilegus,* identified using PhyloP. For each comparison, enrichment was tested using a one-sided Fisher’s exact test. The background was defined as the set of genes eligible for the corresponding *M. spicilegus* candidate analysis.

Synaptic maturation gene sets were from the atlas of proteins detected in postsynaptic density (PSD) of human cortical neurons in (L. Wang et al. 2023). We analyzed, each in turn, the complete set of proteins detected in human PSD (the entirety of Supplementary Table 2 in (L. Wang et al. 2023)); one module of the latter identified by weighted gene co-expression network analysis, which tended to be expressed more highly in human postsynaptic densities during the perinatal period compared with macaque and mouse and was implicated in slower human PSD maturation (the turquoise module from Supplementary Table 2 in (L. Wang et al. 2023)); and members of the latter annotated in the Rho signaling pathway (Supplementary Table 4b-c in (L. Wang et al. 2023)).

The autism-associated gene set comprised 306 genes in which copy number variation was associated with autism from the SPARK autism project (SPARK 2025).

Gene sets related to cichlid bower-building behavior were from single-nucleus transcriptional profiling of courting male fish from (Johnson et al. 2023). We analyzed, each in turn, sets of genes whose expression was associated with building behavior, quivering behavior, or gonadosomatic index (the union of genes that had emerged as markers of clusters from primary, i.e. low-resolution, and secondary, *i.e.* higher-resolution, snRNA-seq and were correlated with the respective behavior in Supplementary Data 7 in (Johnson et al. 2023)).

## Data availability

The data underlying this article are available in the article and its online supplementary material. Publicly available genomic, transcriptomic, regulatory, and gene expression datasets used in this study are available from their original repositories, with accession numbers and source information provided in the Methods and accompanying data manifest in the github repository. Code used for data processing, statistical analyses, and figure generation is available at https://github.com/maracbaylis/MusSpicilegusGenomicSignaturesOfSelection_paper_code. Additional intermediate files are available from the corresponding author upon reasonable request.

## Acknowledgements

We thank Francois Bonhomme for providing stocks of *M. spicilegus*, and Annaliese Beery, George Bentley, Barak Cohen, Lisa Couper, Emily Dennis, Eric Enbody, Lance Kriegsfeld, Claire LeBlanc, George Prounis, Michael Nachman, and members of the Brem and Wilbrecht labs for helpful discussions.

## CRediT authorship contribution statement

**Mara Baylis:** Conceptualization, Methodology, Formal Analysis, Investigation, Data Curation, Visualization, Writing – Original Draft, Writing – Review & Editing. **Kayleigh Fort:** Formal Analysis, Data Curation, Writing – Review & Editing. **Samantha Jackson:** Validation, Writing – Review & Editing. **Linda Wilbrecht:** Conceptualization, Supervision, Resources, Writing – Review & Editing. **Rachel B. Brem:** Conceptualization, Methodology, Supervision, Project Administration, Funding Acquisition, Writing – Review & Editing.

